# Prebiotic proanthocyanidins inhibit bile reflux-induced esophageal adenocarcinoma through reshaping the gut microbiome and esophageal metabolome

**DOI:** 10.1101/2023.08.22.554315

**Authors:** Katherine M. Weh, Connor L. Howard, Yun Zhang, Bridget A. Tripp, Jennifer L. Clarke, Amy B. Howell, Joel H. Rubenstein, Julian A. Abrams, Maria Westerhoff, Laura A. Kresty

**Affiliations:** Department of Surgery, Section of Thoracic Surgery, University of Michigan, Ann Arbor, Michigan, USA; Rogel Comprehensive Cancer Center, University of Michigan, Ann Arbor, Michigan, USA; Department of Electrical and Computer Engineering, Quantitative Life Sciences Initiative, University of Nebraska-Lincoln, Lincoln, Nebraska, USA; Department of Statistics, Department of Food Science Technology, Quantitative Life Sciences Initiative, University of Nebraska-Lincoln, Lincoln, Nebraska, USA; Department of Plant Pathology and Biology, Marucci Center for Blueberry and Cranberry Research, Rutgers University, Chatsworth, New Jersey, USA; Division of Gastroenterology, Department of Internal Medicine, University of Michigan, Ann Arbor, Michigan, USA; LTC Charles S. Kettles Veterans Affairs Medical Center, Ann Arbor, Michigan, USA; Department of Medicine, Columbia University Irving Medical Center, New York, New York, USA; Department of Pathology, University of Michigan, Ann Arbor, Michigan, USA

## Abstract

The gut and local esophageal microbiome progressively shift from healthy commensal bacteria to inflammatory-linked pathogenic bacteria in patients with gastroesophageal reflux disease, Barrett’s esophagus and esophageal adenocarcinoma (EAC). However, mechanisms by which microbial communities and metabolites contribute to reflux-driven EAC remain incompletely understood and challenging to target. Herein, we utilized a rat reflux-induced EAC model to investigate targeting the gut microbiome-esophageal metabolome axis with cranberry proanthocyanidins (C-PAC) to inhibit EAC progression. Sprague Dawley rats, with or without reflux-induction received water or C-PAC *ad libitum* (700 µg/rat/day) for 25 or 40 weeks. C-PAC exerted prebiotic activity abrogating reflux-induced dysbiosis, and mitigating bile acid metabolism and transport, culminating in significant inhibition of EAC through TLR/NF-κB/P53 signaling cascades. At the species level, C-PAC mitigated reflux-induced pathogenic bacteria *(Clostridium perfringens, Escherichia coli,* and *Proteus mirabilis).* C-PAC specifically reversed reflux-induced bacterial, inflammatory and immune-implicated proteins and genes including *Ccl4, Cd14, Crp, Cxcl1, Il6, Il1β, Lbp, Lcn2, Myd88, Nfkb1, Tlr2 and Tlr4* aligning with changes in human EAC progression, as confirmed through public databases. C-PAC is a safe promising dietary constituent that may be utilized alone or potentially as an adjuvant to current therapies to prevent EAC progression through ameliorating reflux-induced dysbiosis, inflammation and cellular damage.

## Introduction

Esophageal adenocarcinoma (EAC) represents a growing health problem characterized by markedly increased incidence in the last 50 years, substantial morbidity and high mortality (1). The only known precursor lesion to EAC is Barrett’s esophagus (BE), a metaplastic protective adaptation to chronic reflux of injurious bile salts and acidic gastric contents, known as gastroesophageal reflux disease (GERD) (1, 2). Acid-reducing proton pump inhibitors (PPIs) are the mainstay of treatment for GERD, but the complete response rate is only about 50% (3, 4). In turn, widespread use of PPIs has not translated to meaningful declines in EAC or improved survival and overall results across studies are inconsistent (5–7). The long-term safety of PPIs has also been questioned, with potential adverse events including reduced gut microbiome health and increased enteric infections (*Clostridium difficile*), deficiencies of micronutrients and renal insufficiency (8–11). Microbial communities progressively shift from healthy commensal bacteria to inflammatory-linked pathogenic bacteria (*Staphylococcus, Streptococcus, Enterococcus* and *Escherichia*) in GERD, BE and EAC (12–18). In the context of microbial dysbiosis pathogenic bacteria are documented to produce DNA-damaging toxins including lipopolysaccharide (LPS) and colibactin as well as proinflammatory metabolites with cancer-promoting effects (19–23). However, mechanisms by which alterations of the gut microbiome-esophageal metabolome axis contribute to EAC progression remain poorly defined and in turn difficult to successfully target. Beyond dysbiosis, reflux components are well documented to induce DNA damage in esophageal cells and to stimulate cytokine-mediated inflammation further contributing toward epithelial injury, immune cell migration, genetic instability and cancer progression (21, 24). Repeated exposure to refluxate also leads to diminished defense mechanisms as evidenced by esophageal barrier dysfunction and reduced levels of esophageal DNA repair enzymes, inferring compromised repair capacity in reflux-exposed esophageal epithelium (21, 25, 26). Reflux-induced mucosal damage coupled with esophageal barrier dysfunction allows for increased interaction between pathogenic bacteria and the epithelium further promoting esophageal injury and risk for EAC progression. Thus, our goal is to identify a safe efficacious agent, for use alone or as an adjuvant, to inhibit reflux-induced EAC by targeting the major drivers and sequelae associated with GERD and BE. Prognosis following a diagnosis of EAC remains poor as evidenced by <20% 5-year survival (27), supporting the urgent need for improved preventive strategies.

Cranberry is a fruit rich in polyphenols, including proanthocyanidins (C-PAC) that exert potent anti-inflammatory and antibacterial activities (28–33) in human trials (31–36) and anti-cancer effects in multiple preclinical models, including EAC xenografts (37–39). Historically, C-PAC was considered to have poor bioavailability and remained underexplored for anti-cancer effects, particularly in vivo. Recently interest in C-PAC was renewed with increased understanding of absorption, digestion and the salient role gut microbes play in generating biologically active metabolites (40–42). Herein, we evaluated the cancer-inhibitory mechanisms of C-PAC utilizing a translationally relevant rat model of reflux-induced EAC. Key findings show that C-PAC exerts prebiotic activity abrogating reflux-induced dysbiosis as evidenced by increasing anti-inflammatory and butyrate-producing bacteria (*Lactobacillus* and *Allobaculum*) while decreasing proinflammatory LPS-producing Gram-negative bacteria (*Escherichia coli*), and mitigating reflux-induced bile acid (BA) metabolism and transport, culminating in inhibition of high-grade dysplasia and EAC through inhibition of Toll-like receptor (TLR)/NF-κB/P53 signaling.

## Results

### Dose range-finding study and C-PAC safety

Rats consuming C-PAC in the drinking water at 250, 500 and 700 µg/rat/day for 6 weeks experienced no adverse effects in terms of body weight, food or water consumption compared to water controls (no significant differences based on repeated measures ANOVA, Supplemental Figure 1A). Additionally, all organ pathology was normal and comprehensive serology measures (Supplemental Figure 1B) did not differ between rats consuming 700 µg/rat/day C-PAC and those receiving water alone.

### C-PAC inhibits progression of reflux-induced EAC

Based on dose-range finding study results, consideration toward the published literature (43, 44) and behaviorally achievable consumption levels, C-PAC was evaluated at a concentration targeting 700 µg/rat/day in the long-term bioassay resulting in actual average delivery of 690 µg/rat/day. As depicted in Figure 1A, following reflux-inducing surgery, animals received treatment of either water alone or C-PAC *ad libitum* in the drinking water. Rats were sacrificed at 25 weeks for histopathology evaluation and spectral imaging (45). At the final 40-week time-point evaluations included histopathology, metabolomics, microbiome profiles and molecular analyses. As shown in Figure 1B, reflux significantly induces HGD and progression to EAC, while reducing the percent of normal-appearing esophageal fields over time. C-PAC effectively mitigated the deleterious impact of reflux as evident by preservation of normal epithelial fields and inhibition of HGD and EAC, most evident at 40 weeks. Compared with reflux alone, the C-PAC+reflux group mitigated EAC formation by 93.6% and 82.4% at 25 and 40 weeks, respectively. C-PAC reduced HGD by about 60% at both time-points. At 25 and 40 weeks, C-PAC+reflux significantly maintained fields of normal epithelium (69.65 and 57.14%), relative to reflux alone (38.83 and 8.01%), supporting that C-PAC potently mitigates refluxant, subsequent tissue damage and progression to EAC.

**Figure 1.**
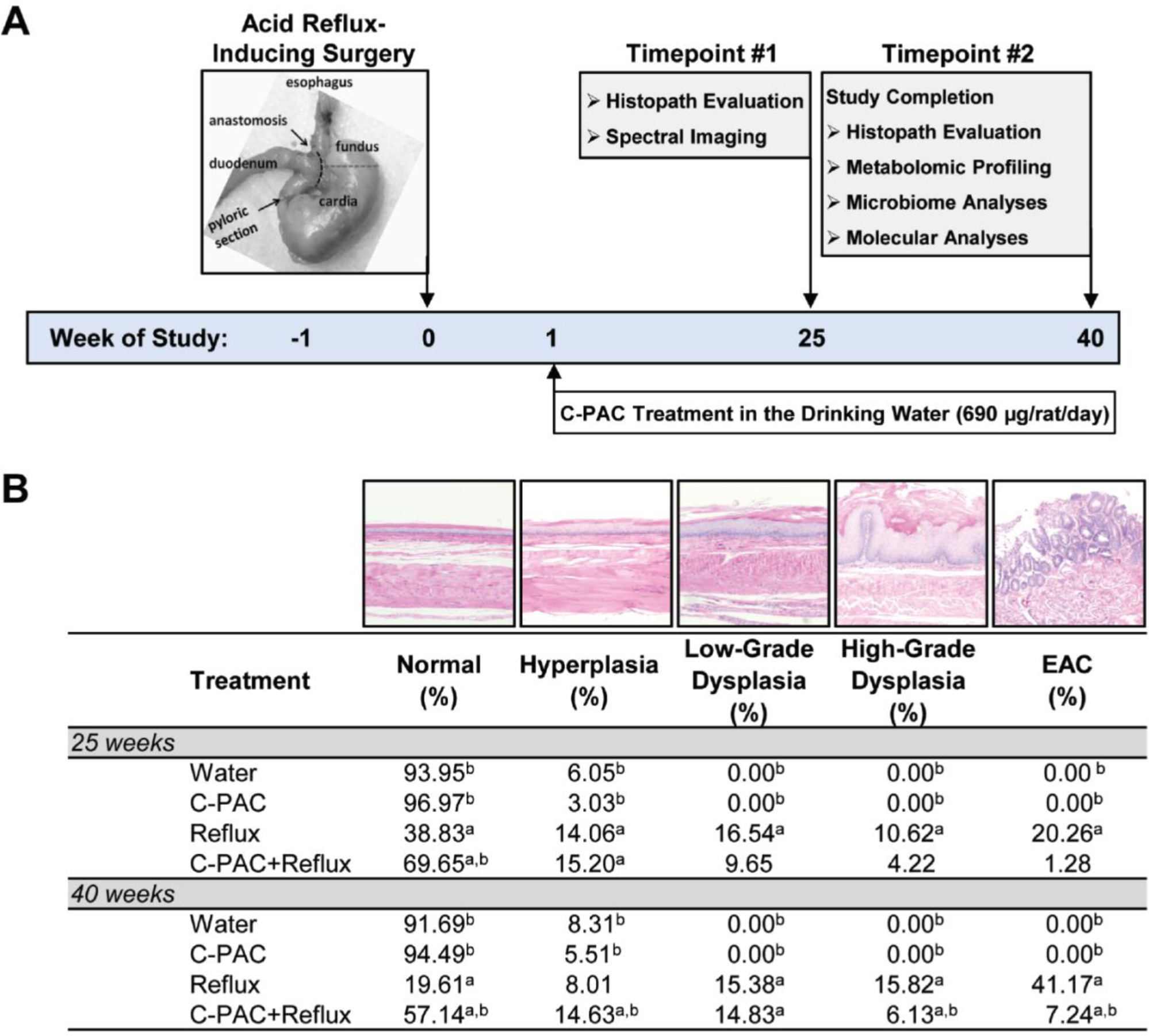
C-PAC inhibits cancer progression in a rat model of reflux-induced EAC. (A) At baseline (week 0) esophagogastroduodenal anastomosis was performed to induce acid-reflux in designated treatment groups. One week later, animals were provided either water or C-PAC (690 µg/rat/day) and AIN93M chow *ad libitum*. At 25 weeks, animals were sacrificed to perform spectral imaging and characterize esophageal histopathology. The study was completed at 40 weeks followed by histopathology evaluation, esophageal metabolomics, gut microbiome profiling and esophageal molecular analyses. (B) Histopathological evaluation of esophagi (n=6 −8 animals per treatment group) were characterized at week 25 and 40 by quantifying the percent of normal, hyperplasia, low-grade dysplasia, high-grade dysplasia and EAC across treatment groups. ^a^Statistically significantly different from water-treated and ^b^Statistically significantly different from Reflux-treated; data are presented as mean values per pathology and analyzed by Students *T*-test (two-sided, *P*≤0.05). C-PAC, cranberry proanthocyanidins; EAC, esophageal adenocarcinoma

C-PAC delivered in the drinking water was safe over the 40-week study duration based on the fact that mean body weight, weight gain and food and water consumption did not differ between the reflux and C-PAC+reflux groups (Supplemental Figure 2A-D, Supplemental Tables 1-4). Body weight differences did emerge between reflux and non-reflux groups due to weight loss during the surgical week from which the rodents never fully rebounded. Importantly, all groups continued to gain weight normally (Table S4) allowing for valid comparisons across treatments.

### C-PAC mitigates reflux-driven proinflammatory gut microbiome changes

To evaluate C-PAC’s capacity to modulate reflux-induced alterations in the gut microbiome, we performed 16S rRNA gene sequencing of fecal pellets collected from negative control or water-treated rats over time and from all treatment groups at week 40. There is reasonable separation of animals by treatment group, with no differences in species richness or alpha diversity, yet statistically significant differences are observed in beta diversity supporting differences in microbial communities in reflux compared with C-PAC+reflux groups (Supplemental Figure 3). Figure 2A shows that consistent with published literature (46), the rat fecal microbiome stabilizes after 2 weeks and the normal microbiome is dominated by the phylum *Firmicutes* followed by *Bacteroidetes* and *Verrucomicrobia*, a composition mimicking normal healthy human esophageal microbiome profiles (12–15). At 40 weeks of study, reflux significantly reduces phylum abundance levels of *Firmicutes* while significantly increasing proinflammatory LPS-linked Gram-negative *Proteobacteria*, *Deferribacteres* and *Bacteroidetes* (Figure 2B)(13, 15, 21). In contrast, C-PAC treatment in the context of reflux results in significant increases in phylum abundance levels of Gram-positive *Firmicutes*, *Tenericutes* and *Actinobacteria* with significant decreases in *Proteobacteria*, as well as marked reductions in *Bacteroidetes* and *Verrucomicrobia* (Figure 2B).

**Figure 2.**
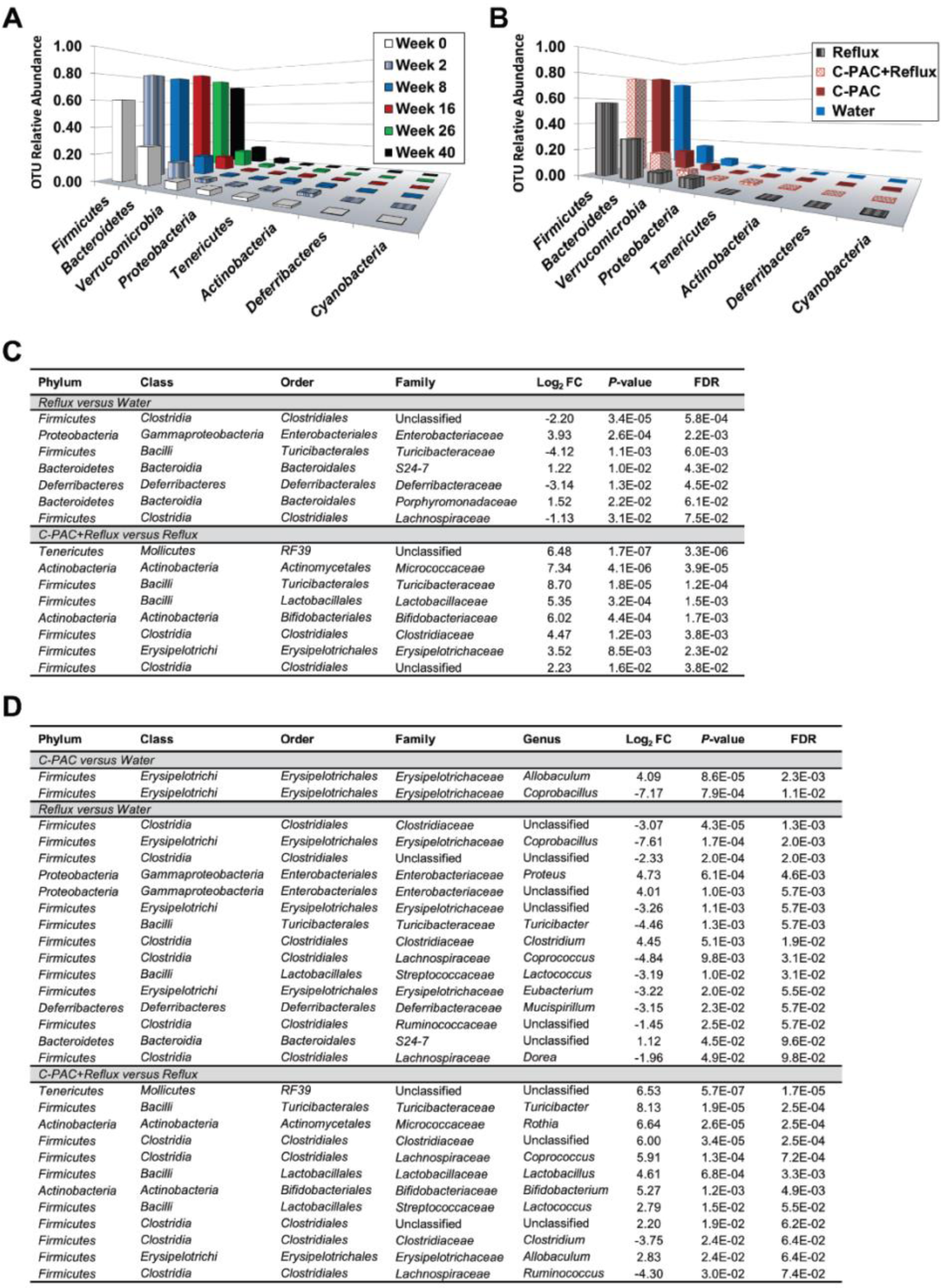
C-PAC ameliorates reflux-induced proinflammatory gut microbiome changes. (A) Gut microbiome profile of water-treated animals from baseline through 40 weeks. Phylum level analysis of 16s rRNA gene sequencing shows stabilization of microbiome profiles after 2 weeks. (B) Gut microbiome profiles at 40 weeks for all treatment groups at the phylum level. (C) Family level changes in the gut microbiome for each treatment group at 40 weeks. (D) Genus level changes in the gut microbiome across treatment groups (40 weeks). Data are presented as Log_2_FC in relative abundance levels with *P*≤0.05 (Robinson and Smyth’s exact test) and FDR≤0.1 (Benjamini-Hochberg correction for false discovery rate). C-PAC, cranberry proanthocyanidins; FC, fold change.

At the family and genus levels, C-PAC in the context of reflux shifts the gut microbiome toward health-associated Gram-positive bacterial populations, increasing beneficial *Firmicutes* and *Actinobacteria* members including *Streptococcaceae, Lactobacillaceae* and *Bifidobacteriaceae* while reducing levels of deleterious *Clostridium* members (Figure 2, C and D)(12). Levels of *Bifidobacteriaceae* were also significantly increased with C-PAC (*P*=0.021; FDR=0.17). At the genus level, C-PAC alone increased levels of *Allobaculum* and *Coprobacillus* (Figure 2D). Conversely, family and genus levels of Gram-negative *Enterobacteriaceae* and *Proteus* were significantly increased with reflux.

Next, we evaluated species level frequency changes utilizing the Axiom microbiome array platform. Results (Supplemental Table 5) reveal that reflux increases pathogenic bacteria including *Odoribacter laneus, Streptococcus mutans, Streptococcus parasanguinis, Proteus mirabilis, E. coli, Enterococcus faecalis* and *Lactobacillus sp ASF360* while reducing acid-resistant anti-inflammatory *Lactobacillius johnsonii* (47). C-PAC mitigated many reflux-induced bacterial species level changes. C-PAC also decreases specific viral families in the context of reflux including *Myoviridae* (*P*=0.0065) and *Siphoviridae* (*P*=0.0549, Supplemental Table 6).

### Esophageal metabolomics reveals novel anti-cancer activities of C-PAC

To characterize metabolic effects of C-PAC in vivo, we conducted untargeted metabolomics on esophageal tissues across treatment groups. A total of 681 compounds were identified, with 319 metabolites significantly different in reflux and 264 metabolites significantly different in C-PAC+reflux (Figure 3A). The PCA plot in Figure 3A shows that esophageal samples separate into well-defined groups by treatment with the greatest variability present with reflux induction. Hierarchical clustering of esophageal samples further supports strong clustering by treatment group (Figure 3B). Next, a biochemical importance plot (Figure 3C) was generated through random forest analysis revealing super pathways and specific metabolites differentiating between treatment groups, with a predictive accuracy of 94%. Top identified super pathways were dominated by amino acids and lipids. Specific biochemicals were linked to redox homeostasis, bacterial pathogenesis and inflammation, including gamma-glutamyl amino acids, lysine, methionine, cytosine, cysteine, glycine as well as numerous sphingomyelins, the proinflammatory lipid prostaglandin E2 and the primary BA chenodeoxycholate (Figure 3C).

**Figure 3.**
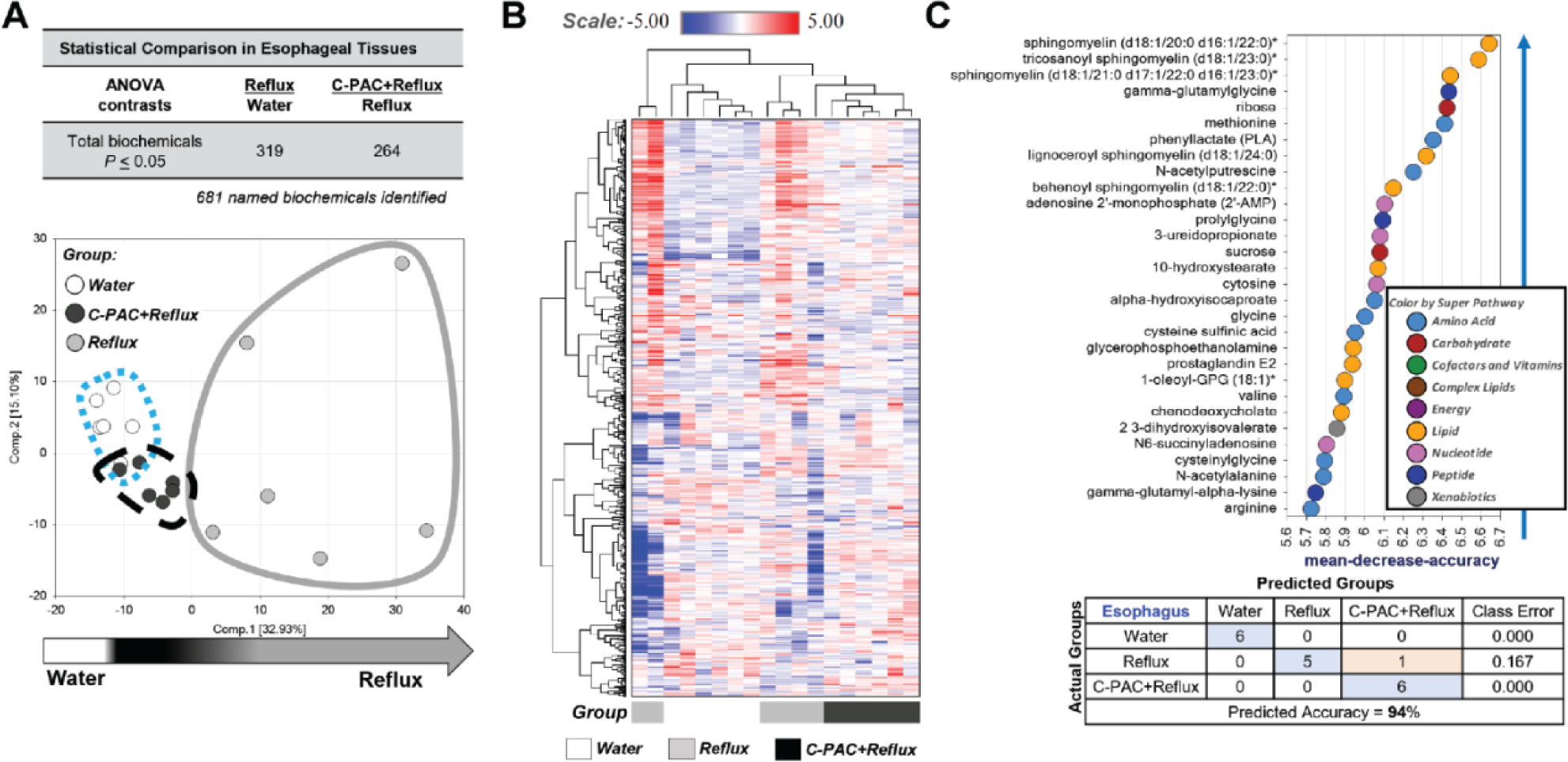
Untargeted metabolomic profiling of esophagi following treatment with C-PAC in reflux-induced EAC. (A) Untargeted metabolomics (n=6 animals per treatment group) revealed 681 named biochemicals identified, with 319 and 264 significantly altered with Reflux or C-PAC in the context of Reflux, respectively. Principal component analysis supported variation between individual samples and treatment groups. (B) Hierarchical clustering of esophageal samples based on named metabolites showed strong clustering based on treatment groups. (C) Biochemical importance plot results identified specific metabolites in super pathways of increasing importance with random forest classification analysis providing a predictive accuracy of 94%. One-way ANOVA was used to identify significant biochemicals with a *P*≤0.05. C-PAC, cranberry proanthocyanidins; EAC, esophageal adenocarcinoma.

Next, significantly modulated metabolites were evaluated based on treatment group comparisons. The Venn diagram in Figure 4A shows that of the 319 metabolites significantly altered by reflux, C-PAC directly reverses 62.7% or 200 metabolites. Top identified pathway maps and metabolic networks derived from the 200 C-PAC mitigated metabolites are summarized in Figure 4B and 4C and Supplemental Tables 7-10. These results align closely with the biochemical importance plot results supporting that reflux up-regulates amino acid metabolism as well as aminoacyl-tRNA biosynthesis and myeloid-derived suppressor and M2 macrophages; all of which are directly reversed by C-PAC. Similarly, Metabolic network analysis (Figure 4C) reveals that C-PAC reverses reflux-induced amino acid metabolism and transport as it relates to glutamate, serine and proline, as well as glycosphingolipid metabolism. Metabolic networks including carnitine, ceramide and phosphocholine pathways were significantly down-regulated by reflux which was directly reversed by C-PAC. As described in Supplemental Table 11, C-PAC mitigates up-regulated reflux-induced Process Networks of translation and vesicle transport. Interestingly, leptin signaling and BA transport are significantly down-regulated by reflux and directly reversed by C-PAC (Supplemental Table 12).

**Figure 4.**
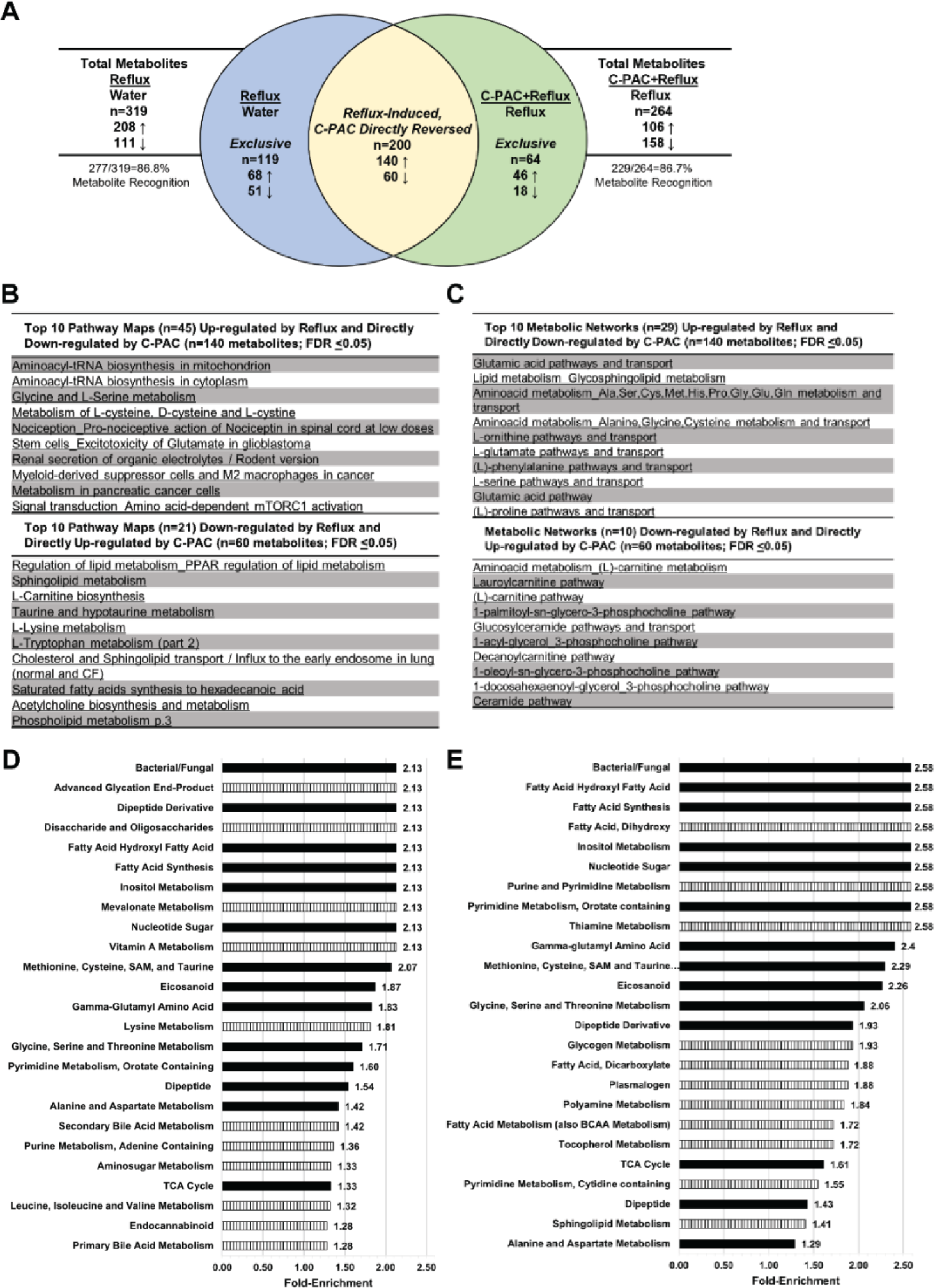
C-PAC mitigates reflux-induced esophageal metabolite dysregulation. (A) Venn diagram shows the total number of significant (*P*≤0.05) metabolites in Reflux vs Water (n=319) and C-PAC+Reflux vs Reflux (n=264). Metabolite recognition in Metacore was 87% in both comparisons. The blue and green sections represent metabolites exclusive to Reflux vs Water (n=119) and C-PAC+Reflux vs Reflux (n=64), respectively. The yellow section represents the metabolites induced by Reflux and directly reversed by C-PAC (n=200). Metabolite directionality is noted as up-regulated (↑) or down-regulated (↓). (B) The top 10 pathway maps for metabolites induced by Reflux and directly reversed by C-PAC in the context of Reflux. Tables S9 and S10 provide complete lists of significant pathway maps identified. (C) The top 10 metabolic networks induced by Reflux and directly reversed by C-PAC in the context of Reflux. Tables S11 and S12 provide complete results of significant metabolic networks detected. Pathway set enrichment analysis performed in Metabolync for (D) Reflux vs Water and (E) C-PAC+Reflux vs Reflux where black bars are shared between comparisons and vertical striped bars are exclusive to the comparison in each panel. Significant pathway maps and metabolic networks are based on an FDR≤0.05. C-PAC, cranberry proanthocyanidins.

Finally, pathway set enrichment analysis via Metabolync identified the top enriched pathways by treatment group as shown in Figure 4, D and E. With reflux, 47 total pathways were significantly enriched with Bacterial/Fungal, Advanced Glycation End-Product and Dipeptide Derivative as the top pathways. Similarly, the top significantly enriched pathway in C-PAC+reflux was also Bacterial/Fungal supporting mitigation of this reflux-induced pathway by C-PAC. Additional pathways modulated by C-PAC in the context of reflux include those associated with fatty acids, eicosanoids, amino acid and nucleotide metabolism, gamma-glutamyl amino acids and the TCA cycle.

### C-PAC mitigates reflux-induced changes in primary and secondary bile acids

Esophageal tissue BAs were assessed considering that reflux of bile and acidic stomach contents has a well-documented role in GERD, BE and EAC progression (1, 2). Fundamentally, BA metabolism includes phase I bile acid synthesis generating relatively equal amounts of the two major primary BAs cholate and chenodeoxycholate in humans and in alignment with our rat model data (48). Next, BAs undergo phase II conjugation to glycine or taurine (Figure 5A) which occurs in a 3:1 ratio in humans and again similarly in rats (48). Conjugation prepares BAs for phase III transport, deconjugation and subsequent conversion by bacteria to highly insoluble and toxic secondary BAs (49). Metabolic profiling reveals that reflux significantly increased levels of multiple primary BAs in rat esophagi including cholate (12.6-fold) and chenodeoxycholate (10.3-fold) and their derivatives, with C-PAC mitigating BA induction (Figure 5B). C-PAC also diminished reflux-induced changes in the conjugating amino acids, glycine and taurine. Box plots of select primary and secondary BAs, as well as BA metabolites are shown in Figure 5C. Bacteria have a well-defined role in transformation of primary BAs into secondary BAs (Figure 5D) including deoxycholate and additional injurious secondary BAs which are up-regulated with reflux and mitigated by C-PAC. Additional quantitative analysis of esophageal cholate and taurocholate confirmed induction by reflux (158.5 nM and 240.8 nM, respectively) and mitigation by C-PAC (100.0 nM and 73.7 nM, respectively). Parallel untargeted metabolomic analysis of liver tissues revealed significantly increased levels of cholesterol (1.3-fold), taurobetamuricholate (2.88-fold) and taurodeoxycholate (4.88-fold) by reflux compared to water controls.

**Figure 5.**
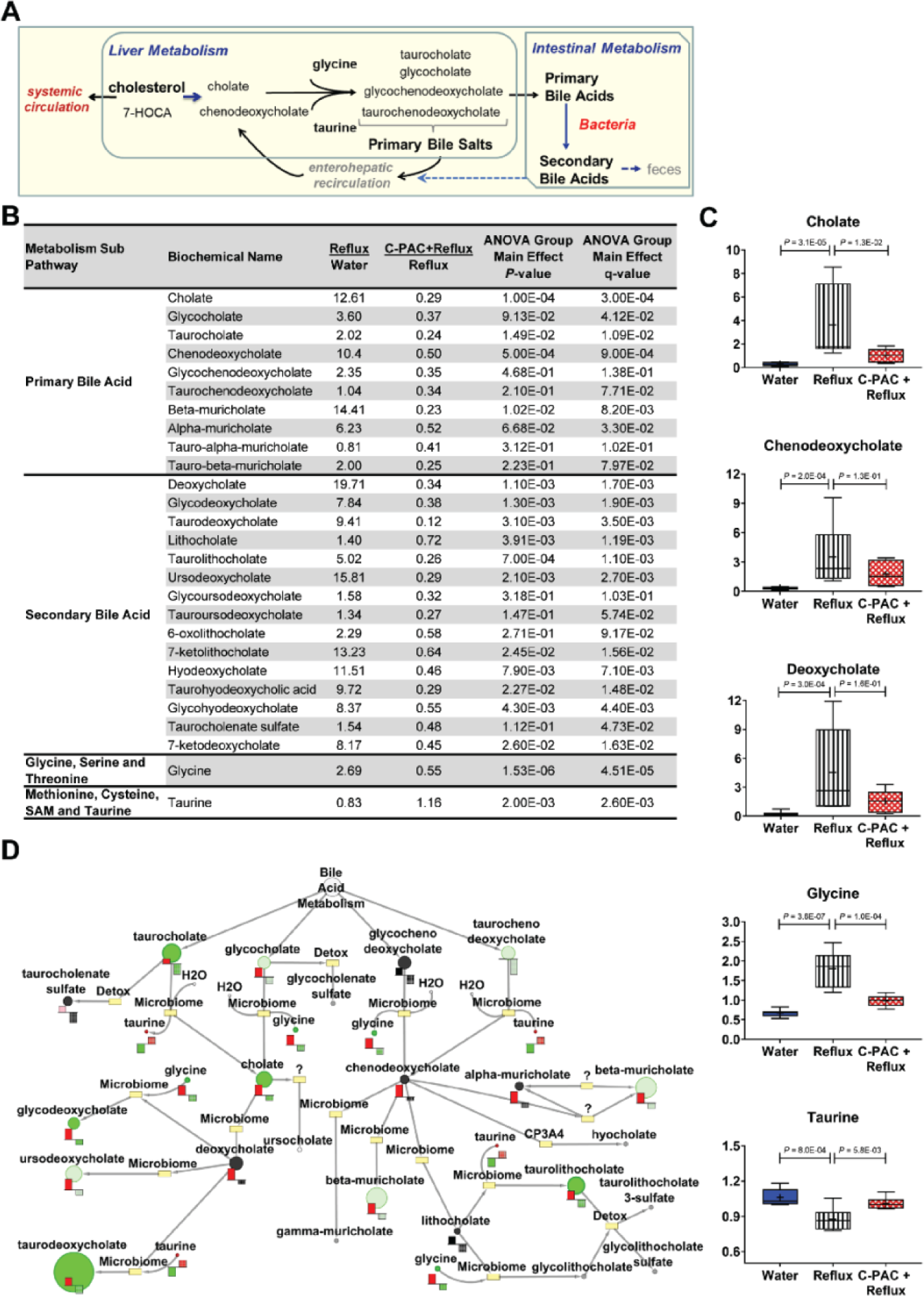
C-PAC mitigates reflux-driven alterations in esophageal bile acids and conjugating amino acids. (A) Summary diagram of BA production and processing. BA metabolism includes phase I BA synthesis in the liver generating relatively equal amounts of the two major primary BAs cholate and chenodeoxycholate from cholesterol in humans and in alignment with our rat model data. Next, BAs undergo phase II conjugation to glycine or taurine in preparation for phase III transport, deconjugation and subsequent conversion by bacteria to highly insoluble and toxic secondary BAs. During gastrointestinal transit most BAs reenter enterohepatic recirculation, leaving only about 5% of BAs to pass through the colon and be excreted in the feces. (B) Primary and secondary BAs, as well as conjugating amino acids with relative fold change levels in Reflux vs Water or C-PAC+Reflux vs Reflux comparisons. Group main effects were determined by one-way ANOVA with Storey’s correction for false discovery rate (q-value). (C) Representative box and whisker plots for select metabolites with data presented as maximal and minimal values (whiskers) and group mean (line); Water (blue), Reflux (white with vertical black stripes) and C-PAC+Reflux (red mesh). *P*-values are reported based on ANOVA for each comparison. (D) BA metabolism pathway generated in Metabolync displaying significantly dysregulated BAs (*P*≤0.05) with circle size indicating relative metabolite level. Levels in bar plots for Reflux vs Water shown in red and C-PAC+Reflux vs Reflux shown in green. C-PAC, cranberry proanthocyanidins; BA, bile acid.

### C-PAC modulates a comprehensive battery of reflux-induced bacterial, inflammatory and immune-related genes and proteins modified in human EAC progression

To further investigate mechanisms by which C-PAC inhibits EAC in vivo, esophageal gene expression analysis was performed targeting a panel of bacterial, inflammatory, and immune-related genes (Table 1). In total 47 genes are significantly dysregulated by either reflux or C-PAC+reflux. Reflux up-regulated markers associated with proinflammatory Gram-negative bacteria (*Cd14, Myd88, Tlr2, Tlr4, Tlr9, Lbp, Nlrp1a, Nlrp3*), activation of NF-κB signaling (*Bcl10, Nfkb1, Nfkbia, Mapk14, Rela, Traf6*), immune recruiting and proinflammatory cytokines linked to BE progression (*Il1b, Il6, Il18, Ccl3, Ccl4, Ccl5, Cxcl1*) with C-PAC mitigating identified changes.

**Table 1.**
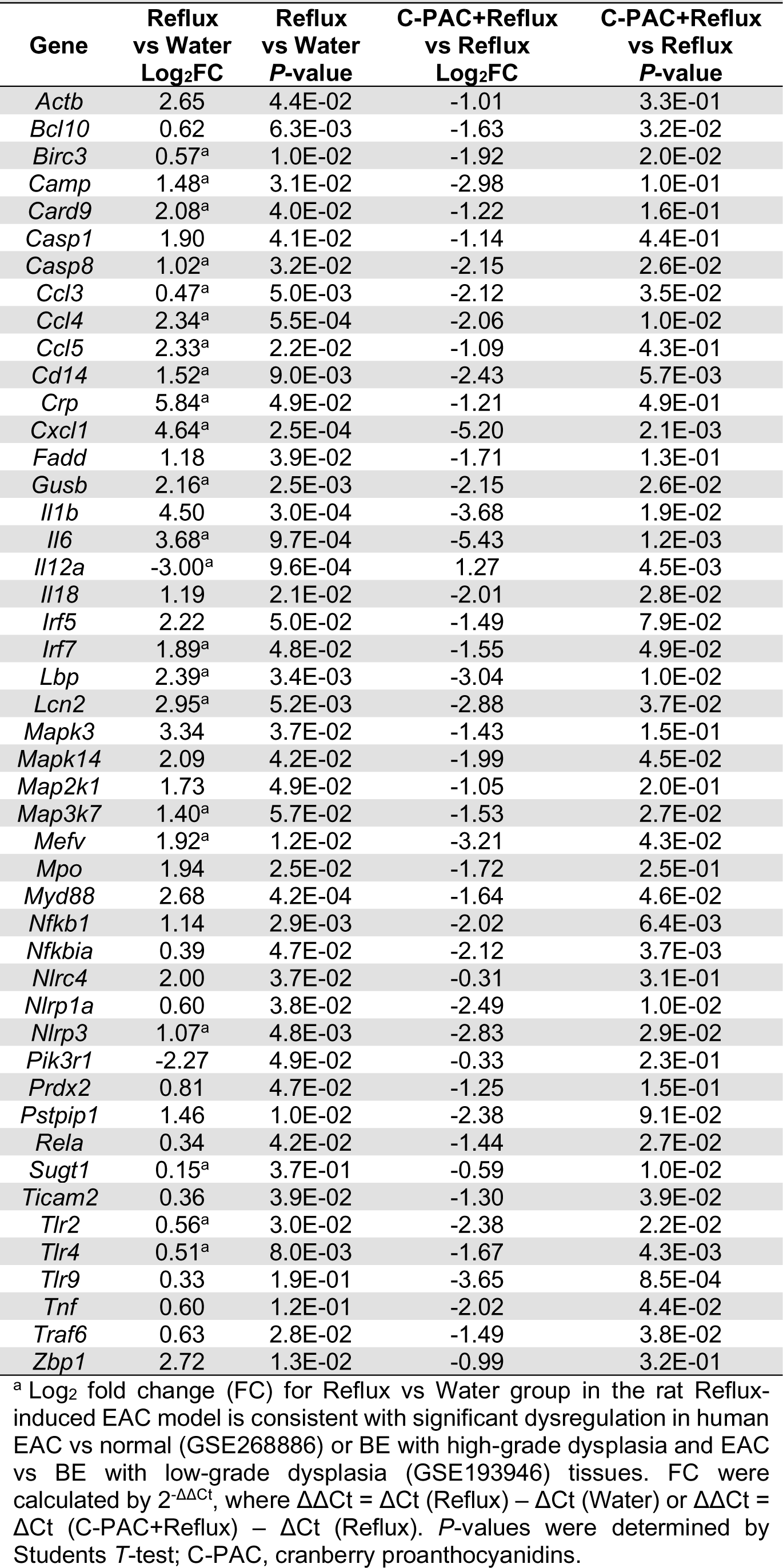
C-PAC mitigates reflux-induced dysregulation of rat esophageal antibacterial genes with direct relevance to human EAC.

To better understand the translational relevance of our results we queried two human data sets for expression changes between histologically normal human esophageal tissues versus EAC (GSE26886), as well as BE tissues with LGD versus BE with HGD and EAC (GSE193946). Of the 47 genes significantly dysregulated in the rat reflux-induced EAC model, 22 markers or 47% were significantly changed and moving in the same direction as in humans shown in Table 1.

In addition to mitigating reflux-induced changes, C-PAC significantly altered expression of antimicrobial markers in normal non-reflux exposed esophagi (Supplemental Table 13), suggesting C-PAC exerts an early protective role preceding bile insult or altered pathology. In normal esophagi, C-PAC altered expression of numerous markers also identified as dysregulated in reflux (Table 1) including *Aps, Bcl10, Camp, Casp1, Ccl4, Ccl5, Cxcl1*, *Il6, Il12a, Irf7, Nfkbia, Prdx2, Sugt1,* and *Ticam2.* Only 5 markers were uniquely altered by C-PAC in normal esophagi including *Ifnb1, Lyz2, Nod2, Prtn3,* and *Pycard*.

In alignment with gene expression results, multiple bacterial- and inflammatory-linked proteins were induced by reflux and mitigated by C-PAC (Figure 6). Increased levels of NF-κB1, TLR3, CD44, COX-2, MyD88, IL-1β and IL-8 are observed with reflux and down-regulated by C-PAC. Consistent with mutations in *TP53* arising with progression to human BE and EAC (50), reflux increased mutant P53 levels with mitigation by C-PAC (51, 52). Reflux increased MAPK signaling through phosphorylation SAPK/JNK^T182/Y185^, ERK1/2^T202/Y204^ and P38^T180/Y182^, while C-PAC mitigated MAPK signaling through reduced phosphorylation of SAPK/JNK and ERK1/2. Finally, consistent with human studies (53), increased levels of cell replication (PCNA) and retinoic acid receptor signaling (RXRγ) were observed with reflux and mitigated by C-PAC. Nuclear receptors, including retinoid family members regulate host-microbiome crosstalk (54).

**Figure 6.**
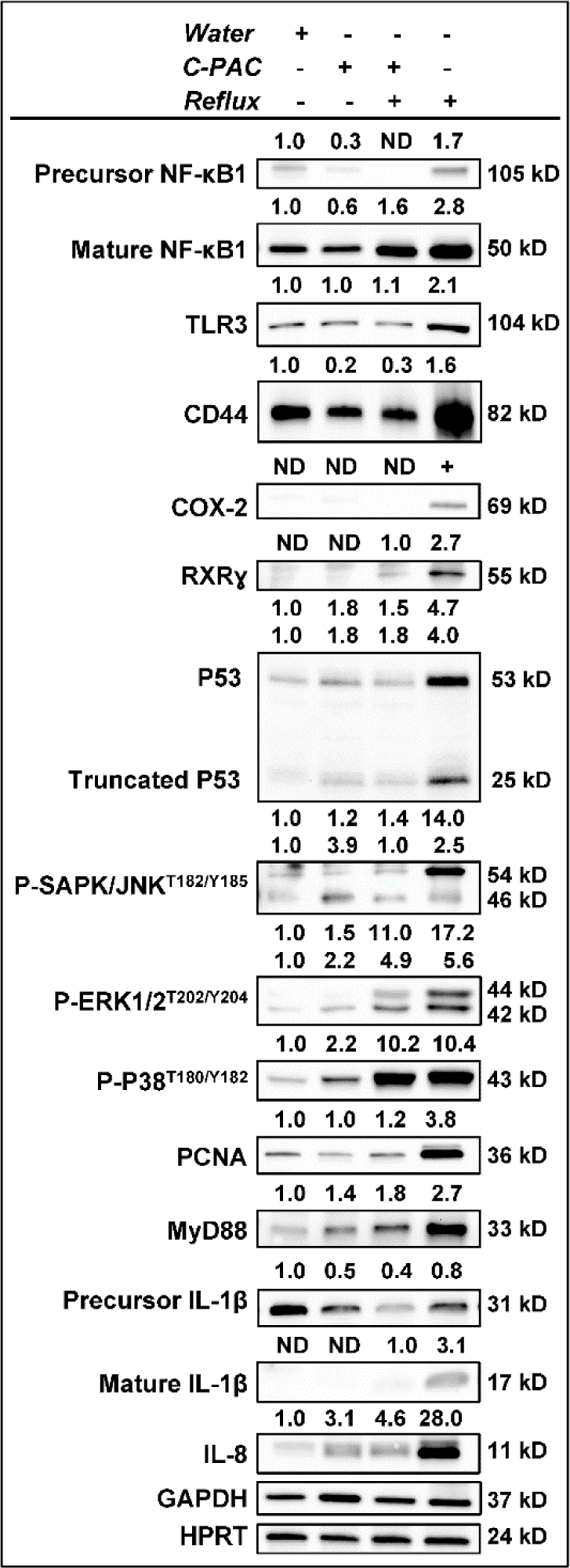
C-PAC mitigates bacterial, inflammatory and immune-related markers dysregulated by reflux. At 40 weeks of study Western blot analysis of esophageal lysates was performed using commercially available antibodies to proteins of interest as described in Table S2. The plus (+) sign denotes treatment group. Expression values were normalized to the appropriate loading control, GAPDH, HPRT or Histone H3 and fold change from Water was calculated using ImageLab. Bands with no detectable expression are denoted as ND. C-PAC, cranberry proanthocyanidins.

### Integration of metabolites and gene expression data reveals networks significantly altered by reflux and mitigated by C-PAC

Given alterations in microbiome profiles, bacterial metabolites and inflammatory signaling cascades with reflux, we utilized Metacore integration to investigate metabolite-gene networks based on significantly dysregulated metabolites in reflux vs water (n=319) and C-PAC+reflux vs reflux (n=264), coupled with significantly altered antibacterial genes (n=47). Of note, Network 2 focuses on BA signaling through Taurocholic acid, Networks 4-6 and 8-10 describe NF*-*κB signaling; whereas, Networks 4 and 7 highlight signaling through TLRs 2, 4 and 9 (Supplemental Table 14), further supporting a strong role for the microbiota in bile metabolism and inflammation. Networks 2 and 5 pictorially depict linkages between metabolite and gene signaling, including the BAs taurodeoxycholate and glycodeoxycholate, as well as the *c-Myc/STAT3/IL-6/* central signaling node, respectively (Supplemental Figure 4).

### Functional microbiome predictions using PICRUSt highlights the importance of transport, secondary metabolism, biofilms and lipid biosynthesis for C-PAC’s anti-cancer activity in vivo

To infer functional consequences of gut microbiome changes in reflux-induced EAC and mitigation by C-PAC, we utilized Phylogenetic Investigation of Communities by Reconstruction of Unobserved States (PICRUSt). KEGG Orthology (KO) and pathways were characterized at classification Levels 2-3 (Table 2). Predicted significant (*P*<0.05, FDR<0.05) functional pathways at Level 2 related to metabolism, specifically Glycan biosynthesis and metabolism, as well as Lipid metabolism. Level 3 predicted functional pathways significantly changed across treatments (ANOVA *P*≤0.05) including Biosynthesis of ansamycins, Bacterial invasion of epithelial cells, Pathogenic *Escherichia coli* infection, ABC transporters and Glutathione metabolism in alignment with other results herein.

**Table 2.** KEGG Ontologies highlight functional microbiome alterations in C-PAC inhibition of reflux-induced EAC.

|  |  |  |  |  |
| --- | --- | --- | --- | --- |
| **Classified to Level 2** |  |  |  |  |
| **Level 1** | **Level 2** |  | **P-value** | **P-value (corrected)** |
| Human Disease | Immune System Diseases |  | 1.3E-03 | 2.4E-02 |
| Metabolism | Glycan biosynthesis and metabolism |  | 7.1E-04 | 2.4E-02 |
| Human Diseases | Infectious Diseases |  | 2.2E-03 | 2.7E-02 |
| Poorly Characterized | Function unknown |  | 3.3E-03 | 3.0E-02 |
| Metabolism | Lipid Metabolism |  | 9.2E-03 | 6.7E-02 |
| Organismal Systems | Excretory System |  | 1.7E-02 | 1.0E-01 |
| Poorly Characterized | General function prediction only |  | 3.3E-02 | 1.7E-01 |
| Genetic Information Processing | Replication, recombination and repair proteins |  | 4.6E-02 | 2.1E-01 |
| **Classified to Level 3** |  |  |  |  |
| **Level 1** | **Level 2** | **Level 3** | **P-value** | **P-value (corrected)** |
| Metabolism | Metabolism of Terpenoids and Polyketides | Biosynthesis of ansamycins | 5.8E-04 | 1.6E-01 |
| Metabolism | Lipid Metabolism | Alpha-Linolenic acid metabolism | 1.5E-03 | 2.1E-01 |
| Human Diseases | Metabolic Diseases | Type I diabetes mellitus | 4.9E-03 | 3.6E-01 |
| Human Diseases | Infectious Diseases | Bacterial invasion of epithelial cells | 6.7E-03 | 3.6E-01 |
| Human Diseases | Cancers | Bladder cancer | 7.0E-03 | 3.6E-01 |
| Human Diseases | Infectious Diseases | Shigellosis | 9.0E-03 | 3.6E-01 |
| Metabolism | Xenobiotics Biodegradation and Metabolism | Bisphenol degradation | 7.8E-03 | 3.6E-01 |
| Genetic Information Processing | Folding, Sorting and Degradation | Chaperones and folding catalysts | 1.4E-02 | 3.8E-01 |
| Human Diseases | Infectious Diseases | Pertussis | 1.5E-02 | 3.8E-01 |
| Human Diseases | Infectious Diseases | Pathogenic *Escherichia coli* infection | 1.5E-02 | 3.8E-01 |
| Metabolism | Xenobiotics Biodegradation and Metabolism | 1,1,1-Trichloro-2,2-bis(4-chlorophenyl)ethane (DDT) degradation | 1.5E-02 | 3.8E-01 |
| Organismal Systems | Excretory System | Proximal tubule bicarbonate reclamation | 1.6E-02 | 3.8E-01 |
| Metabolism | Xenobiotics Biodegradation and Metabolism | Toluene degradation | 2.2E-02 | 4.5E-01 |
| Metabolism | Lipid Metabolism | Linoleic acid metabolism | 2.2E-02 | 4.5E-01 |
| Environmental Information Processing | Signaling Molecules and Interaction | Ion channels | 3.8E-02 | 5.5E-01 |
| Environmental Information Processing | Membrane Transport | ABC transporters | 4.3E-02 | 5.5E-01 |
| Genetic Information Processing | Transcription | Transcription factors | 4.2E-02 | 5.5E-01 |
| Metabolism | Metabolism of Terpenoids and Polyketides | Geraniol degradation | 3.5E-02 | 5.5E-01 |
| Metabolism | Energy Metabolism | Photosynthesis proteins | 3.7E-02 | 5.5E-01 |
| Metabolism | Metabolism of Terpenoids and Polyketides | Biosynthesis of vancomycin group antibiotics | 3.9E-02 | 5.5E-01 |
| Metabolism | Metabolism of Other Amino Acids | Glutathione metabolism | 4.0E-02 | 5.5E-01 |
| Metabolism | Energy Metabolism | Photosynthesis | 4.1E-02 | 5.5E-01 |
| Metabolism | Metabolism of Cofactors and Vitamins | Vitamin B6 metabolism | 4.6E-02 | 5.6E-01 |
| Metabolism | Biosynthesis of Other Secondary Metabolites | Stilbenoid, diarylheptanoid and gingerol biosynthesis | 4.8E-02 | 5.6E-01 |

Multiple group statistical analysis performed in STAMP with *P*-values based on ANOVA with Tukey's post-hoc test and Storey's correction for multiple comparisons (*P*-value corrected). KEGG, Kyoto Encyclopedia of Genes and Genomes; C-PAC, cranberry proanthocyanidins; EAC, esophageal adenocarcinoma.

Functional analysis (Figure 7A-H; Supplemental Table 15) identified 125 KOs significantly modulated. Multiple genes linked to transport including *aapJ, uraA, tolA, tolR, lptE, phaA, phaC-G* and *bamA* (Figure 7A, 7E) were significantly up-regulated with reflux and reversed by C-PAC. Consistent with *aapJ* belonging to the bacterial family of ABC transporters and being up-regulated with reflux, we observed increases in the BA transporter ABCB1 and potent mitigation by C-PAC at the protein level (Figure 7A).

**Figure 7.**
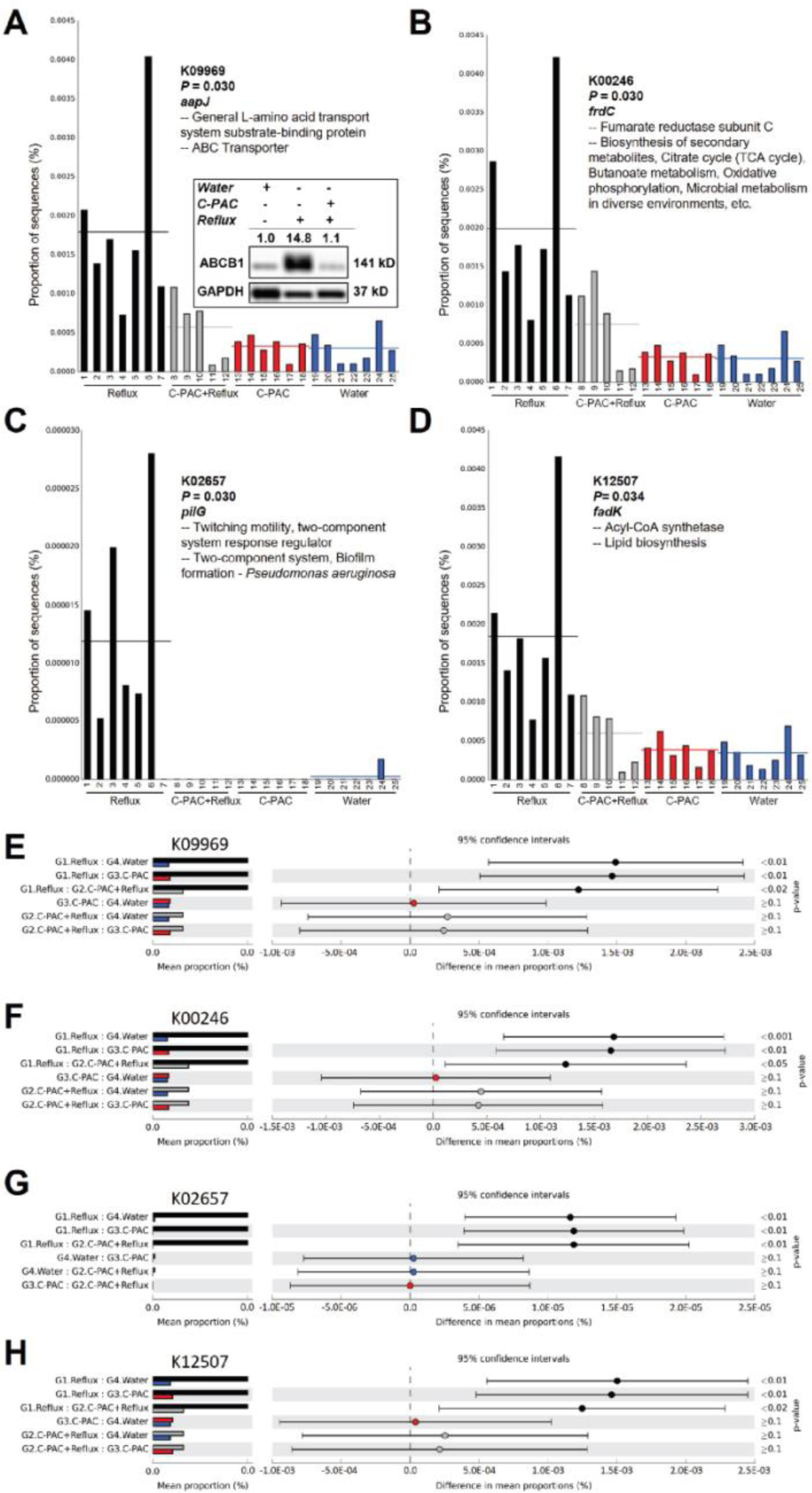
Functional microbiome predictions via PICRUSt reveal novel roles for C-PAC in vivo. 16S rRNA gene sequencing of fecal pellets from animals at 40 weeks was analyzed using PICRUSt and visualized in Statistical Analysis of Metagenomic Profiles (STAMP). Highlighted results by animal and treatment assignment (A-D) and groupwise comparisons based on treatment (E-H) are shown; in (A and E) *aajP*, a general L-amino acid ABC transporter; (B and F) *frdC*, fumarate reductase subunit C; (C and G) *pilG*, twitching motility gene; (D and H) *fadK*, an acyl-CoA synthetase involved in lipid biosynthesis. Treatment groups are denoted by color: Reflux (black), C-PAC+Reflux (gray), C-PAC (red) and Water (blue). Data were analyzed by ANOVA with Tukey’s post-hoc test and Storey’s correction for FDR. The western blot inset in (A) is of the BA ABC transporter ABCB1 in rat esophageal tissues at 40 weeks. The plus (+) sign denotes treatment group. ABCB1 protein levels were normalized to GAPDH and fold change from Water was calculated. C-PAC, cranberry proanthocyanidins; BA, bile acid.

Additionally, multiple genes with roles in bacterial metabolism are significantly dysregulated with reflux and restored by C-PAC (Figure 7, B and F) including *frdC, glpK, mlycD, fumB, pccA* and *ligA* which are linked to biosynthesis of secondary metabolites, tricarboxylic acid cycle (TCA), butanoate metabolism, oxidative phosphorylation and propanoate metabolism (Supplemental Table 15). Bacterial twitching and biofilm formation occur through *pilG* which is up-regulated with reflux and ablated with C-PAC (Figure 7, C and G). Last, enzymes linked to lipid biosynthesis including *fadK* (Figure 7, D and H) and peptidoglycan biosynthesis as it relates to Glycan biosynthesis and metabolism (*ampD, mltA, pbpG)* were up-regulated with reflux and down-regulated by C-PAC, consistent with observed changes in the gut microbiome and bacterial metabolites.

## Discussion

Our laboratory has previously shown the cancer inhibitory potential of C-PAC utilizing human esophageal normal, BE and EAC cell lines and OE19 EAC tumor xenografts (25, 37–39, 55). The current study extends evaluation of C-PAC to a more translationally relevant reflux-induced EAC model. Importantly, this rat model permits assessment of C-PAC in the context of the main risk factor driving the development of BE and EAC progression (1, 2), namely reflux of bile and acidic gastric contents into the lower esophagus. Oral delivery of C-PAC (690 µg/rat/day) was well tolerated throughout the chemoprevention bioassay and resulted in potent anti-cancer effects when ingested at behaviorally achievable levels (human equivalent of 36 mg/day or 10 oz of 27% cranberry juice). While we and others have shown the cancer inhibitory potential of C-PAC in multiple in vitro models (39), the current study is the first to show C-PAC inhibits reflux-driven EAC in vivo and to decipher mechanisms of inhibition linked to reflux-driven dysbiosis, transport and metabolism of bile, and TLR/NF-κB/TP53 signaling.

The role of the gut microbiota influencing systemic inflammation is well documented, but less is known regarding responses at remote tissue sites including the esophagus. Emerging evidence supports that alterations in microbiota and metabolites are linked to GERD, BE, and EAC progression (12–16, 56). The microbiome among these patients is increasingly dominated by Gram-negative LPS-producing pathogenic bacteria (13–18), as is the microbiome in the rat reflux-induced EAC model. Many proinflammatory family- and species-level bacteria (i.e. *Enterobacteriaceae, Verrucomicrobiaceae, Proteus* sp.) are increased in BE and EAC patients (14) and similarly induced in the rat reflux model with significant mitigation by C-PAC. C-PAC restored reflux-induced gut microbiota alterations to a more healthful, anti-inflammatory state with increased levels of Gram-positive *Firmicutes* phylum members. C-PAC induced expression of health-associated *Lactococcus, Lactobacillus* and *Bifidobacterium*, bacteria frequently administered as probiotics (57). Furthermore, *Allobaculum*, a Gram-positive butyrate producing bacterium (58), was significantly increased by C-PAC alone. Recently, Ma et al. reported two members of the gut virome, *Myoviridae and Siphoviridae,* increased with progression from BE to EAC (59), and we report for the first time that C-PAC decreased the frequency of these viral families in the context of reflux. Virome level changes were postulated to be linked to LPS-induced alterations in the gut microbiome composition resulting in inflammation and EAC progression. Thus, modulation of the gut microbiome and virome by C-PAC is consistent with abrogating inflammatory-linked microbial pathogenesis induced by reflux and contributing to EAC progression. To date, few human studies utilizing cranberries have been completed that profile the microbiome; however, two trials do support that cranberries exert beneficial effects on the microbiome. Two weeks of a single serving of sweetened dried cranberries increased commensal gut bacteria and decreased pathogenic bacteria in healthy subjects (34). Similarly, a short duration randomized controlled trial of 11 healthy subjects fed an animal-based diet linked to dysbiosis, reported that cranberry powder attenuated effects by increasing *Firmicutes* and short-chain fatty acids while decreasing *Bacteroidetes* and, interestingly, reducing deoxycholic and lithocholic secondary BAs, both implicated in reflux (33). These studies provide direct evidence that dietary cranberry products favorably influence the gut microbiome and microbial metabolites in healthy humans however, studies in patients with altered esophageal pathologies are nonexistent.

Considering that microbiota exert pro-carcinogenic effects by influencing host metabolism (19–23), we conducted untargeted metabolomics across treatment groups revealing alterations in microbial, immune and inflammatory-linked esophageal metabolites with reflux and mitigation by C-PAC. Most notable is the impact of C-PAC on reflux-induced BAs, the main exposure contributing to EAC progression (1, 2). C-PAC diminished levels of both primary and bacterial-derived secondary BAs in reflux-exposed esophagi. Favorable metabolic findings were further supported by Pathway map, Metabolic network and enrichment analyses revealing that reflux alters Bacterial/fungal, Fatty Acid Synthesis, Methionine/Cysteine/SAM/Taurine, Gamma-Glutamyl Amino Acid, TCA cycle and Eicosanoid Metabolism, with C-PAC reversing these proinflammatory cancer-linked alterations. Additional specific metabolites up-regulated by reflux and mitigated by C-PAC included glycine, glutamate, glutamine and lysine, all metabolites identified in esophageal tissues and serum from EAC patients (60, 61).

Process network and functional microbiome prediction using PICRUSt provided additional mechanistic insight regarding C-PAC’s prebiotic cancer inhibitory activity. Several transporter KOs were significantly up-regulated by reflux and down-regulated by C-PAC including the ABC transporter K09969 and the BA efflux pump *ABCB1* also known as *MDR1* due to its role in transport of anti-cancer agents (62). In accord with this, Snider et al. published that ABC transporters were enriched in HGD and EAC samples compared to non-dysplastic BE and LGD human esophageal tissues (14). Additional transporters are known to be dysregulated in EAC; however, much of the research has focused on therapeutic resistance, not cancer prevention (63–65). C-PAC also reversed reflux-induced K05501 and K02569 with linkages to antibiotic resistance, as well as K04015 which is associated with nitrate/nitrite reductases are significantly dysregulated in EAC progression (17). It is predictable that agents like C-PAC that normalize transport may have efficacious roles in cancer prevention as well as adjuvant therapy. In parallel with our metabolomic pathway map results, PICRUSt also identified altered Glutathione metabolism by reflux and mitigation by C-PAC. These data align with earlier research by our group revealing that C-PAC protects patient-derived normal primary esophageal cultures against BA-induced damage through induction of Glutathione S-transferase Theta 2 (GSTT2), a phase II detoxification enzyme with postulated roles in esophageal mucosal defense and cancer inhibition (25, 26). Similarly, we also reported that C-PAC reversed reflux-induced loss of GSTT2 while simultaneously decreasing esophageal DNA damage, supporting a protective role for this protein targeting EAC in vivo (26). Thus, PICRUSt functional analysis provides further evidence that reflux induces deleterious microbiome changes that are mitigated by C-PAC, in part through effects on transport and detoxification, contributing to EAC inhibition.

Molecular analyses further characterized C-PAC’s capacity to mitigate reflux-linked modulation of the gut microbiome, microbial metabolites, inflammation and immune signaling. LPS is a toxic bacterial metabolite secreted by Gram-negative bacteria and known to trigger host immune responses through receptors including Toll-family members and LPS-binding protein (LBP), leading to activation of NF-κB and other inflammatory signaling cascades linked to reflux-driven EAC (66). C-PAC modifies signaling associated with bacterial and viral host recognition known to be dysregulated in EAC progression as evidenced by C-PAC modulating reflux-induced effects on Toll-like receptors 2-4 and 9, as well as LBP (21, 67). C-PAC also modulates reflux-induction of CD44, a multifunctional transmembrane glycoprotein that regulates TLR activity and plays a role in barrier function through tight-junction assembly (68, 69). At the transcript and protein level, C-PAC mitigated immune recruiting, proinflammatory cytokines linked to BE progression (24), markers associated with increased Gram-negative bacteria and activation of the NF-κB pathway, which is also known to be induced by specific BAs (21, 70). Moreover, for the first time we show that C-PAC decreased esophageal levels of mutant P53 in the context of reflux in vivo, consistent with our previous in vitro research (37, 38). Mutant P53 levels have been noted in the rat reflux-induced EAC model with accumulation of P53 in nuclei and the cytoplasm of bile-developed columnar-lined epithelium and EAC tissues at 40 weeks (71). The gut microbiome alters oncogenic functions of P53 (72), raising the question of whether *TP53*-linked cancers can be inhibited by microbiome-targeting agents such as C-PAC. Orally delivered C-PAC proved efficacious at inhibiting EAC progression through mitigating reflux-induced inflammation, immune suppression and dysbiosis while reducing levels of mutant P53. Additionally, C-PAC reversed reflux-induced increases in MAPK signaling via SAPK/JNK and ERK1/2 but not P38. These results are consistent with our earlier in vitro findings in BE and EAC cell lines, as well as in OE19 xenografts, following C-PAC treatment (37, 38). The sustained signaling of P38 with C-PAC in the context of reflux combined with reflux-driven increased levels of mutant P53, may be in part due to MAPK and P53 crosstalk (73). Collectively these results highlight the potent effects of C-PAC on EAC inhibition not only through microbial, but metabolic and molecular reprogramming of the reflux-exposed esophagus.

Despite limitations and knowledge gaps, results from the present study hold strong potential for informing future clinical studies targeting GERD, BE and EAC inhibition. Study limitations include that BA interrogation was solely conducted at the tissue level without capacity to assess the total BA pool. Still, in alignment with potential impact on the BA pool and associated systemic effects, reflux-induction significantly elevated liver cholesterol and taurine-conjugated BAs. Microbiome species level changes were also restricted to frequency-based measures in the current study. Furthermore, clinical trials assessing cranberry polyphenols or C-PAC have mainly focused on infections related to the urinary tract, *Helicobacter pylori, Escherichia coli* or select pathogenic microbes (44, 74, 75). In fact, C-PAC was tested in only a single cancer cohort showing that 30 days of oral delivery significantly reduces serum prostate-specific antigen levels (22.5%) in prostate cancer patients (76). In summary, C-PAC represents a safe, efficacious and widely available dietary constituent that acts as a prebiotic to mitigate reflux-induced inflammation and damage in a translationally relevant rat EAC model through modulation of the gut microbiome, microbial metabolite levels and immune signaling cascades requisite for BE progression to EAC (Figure 8). Additional research is warranted to fully characterize effects of C-PAC on specific immune cell populations and barrier function. Future research should also include clinical evaluations of C-PAC in patients with GERD or BE, particularly those with mutant *TP53* or at increased risk for EAC progression based on modeling or the presence of additional risk factors. GERD and BE are the strongest known risk factors for EAC followed by obesity, especially abdominal or visceral obesity (77). A polyphenol-rich cranberry extract protects from diet-induced obesity, visceral obesity, insulin resistance and intestinal inflammation in association with increased *Akkermansia* spp. population in the gut microbiota of mice supporting favorable effects in the context of an additional key risk factor for EAC (78). This aligns with our results revealing Lipid Metabolism as a top functional microbiome alteration linked to C-PAC inhibition of reflux-induced EAC in our rat model. Additionally, a recent randomized placebo-controlled clinical trial reported that 8 weeks of consumption of a cranberry beverage rich in C-PAC lead to a reduction of triacylglycerol and oxidative stress levels in patients with elevated levels of C-reactive protein supporting favorable effects in humans that parallel changes identified in preclinical models (79). Mounting evidence supports that microbial alterations contribute to reflux-induced BE and progression to EAC through key roles in metabolism, inflammatory signaling and mucosal defense. Thus, C-PAC seems especially well suited for targeting BE and EAC, as it mitigates the major risk factors, as well as the biologic and molecular sequelae.

**Figure 8.**
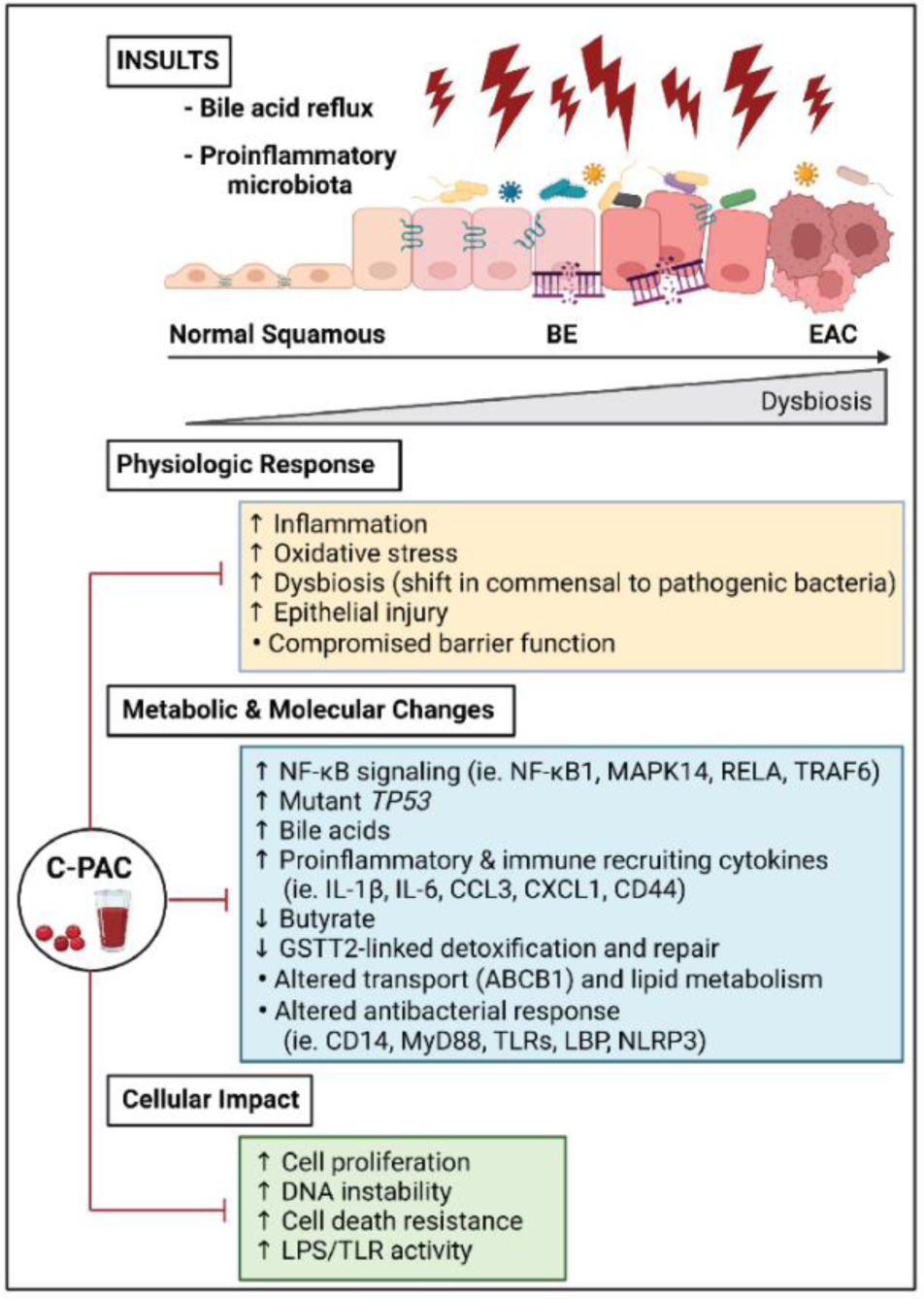
Schematic summary of C-PAC ameliorating reflux-driven alterations in the microbiome-metabolite-host axis supporting EAC inhibition. Chronic reflux of injurious gastric and duodenal contents into the esophagus promotes BE by triggering inflammation, DNA damage, and esophageal barrier dysfunction which in turn allows for increased interaction between pathogenic bacteria and the epithelium further driving esophageal injury and escalating EAC risk. BA refluxate also contributes to gut and esophageal dysbiosis in GERD, BE and EAC patients shifting the microbiome in a proinflammatory direction, as similarly noted in the rat-reflux-induced EAC model with C-PAC potently reversing gut dysbiosis and counteracting metabolic and molecular changes, as well as physiologic and cellular responses as outlined. C-PAC has profound effects on decreasing signaling through NF-κB, levels of mutant *TP53*, primary and secondary BAs, as well as proinflammatory and immune recruiting cytokines while increasing short chain fatty acids (ie. butyrate) and repair linked GSTT2 levels. Restoration of these metabolic and molecular changes by C-PAC resolves reflux-induced BA metabolism and transport dysregulation supporting reestablishment of host antibacterial responses and epithelial barrier function. Finally, C-PAC mitigates reflux-induced increases in cellular proliferation, DNA instability, cell death resistance and LPS/TLR activity. Taken together study results support that C-PAC inhibits reflux-induced high-grade dysplasia and EAC through targeting the major drivers of BE in parallel with restoration of the gut microbiome, reduced inflammation and modulation of cancer-associated TLR/NF-κB/P53 signaling. BA, bile acid.

## Methods

### Cranberry proanthocyanidins source, isolation and preparation

C-PAC was sourced and isolated as previously described (37). Briefly, fresh cranberries (*Vaccinium macrocarpon* Ait.) were collected at the Marucci Center for Blueberry and Cranberry Research (Chatsworth, NJ) and purified C-PAC was isolated from cranberries of the ‘Early Black’ cultivar using solid-phase chromatography according to well established methodology (43, 80–82). Following homogenization in 70% aqueous acetone, the mixture was filtered and pulp discarded. The resulting C-PAC fraction was concentrated under reduced pressure and purified extract isolated using bioassay-directed fractionation. The absence of absorption at 360nm and 450nm confirmed all molecules except the proanthocyanidins were removed. To verify the presence of A-type linkages and proanthocyanidin concentration, ^13^C NMR, electrospray mass spectrometry, matrix-assisted laser desorption/ionization time-of-flight mass spectrometry and acid-catalyzed degradation with phloroglucinol were used, respectively. Purified C-PAC was freeze-dried and stored at −80°C. To prepare C-PAC in the drinking water for study, C-PAC was first dissolved in 0.001% ethanol and vivarium water, sonicated and dissolved to the appropriate final concentration in vivarium water. Fresh C-PAC was replenished at least weekly, as described below.

### Animals

Following a one-week period of acclimation to the facility, 5-7 week old male Sprague-Dawley rats (Crl:SD, strain code 400; Charles River Laboratories, Wilmington, MA) were weighed and randomized into experimental groups as indicated below for the dose-range finding study (n=24) or the long-term chemoprevention bioassay (n=130). Rats were pair-housed in a controlled environment with standard light/dark and temperature conditions and were fed AIN93M purified diet (Dyets Inc., Bethlehem, PA) and water or C-PAC in the drinking water available *ad libitum*. Body weight, food and water consumption were measured at least weekly.

### The rat reflux-induced EAC model

Reflux-induced EAC was created via an esophagogastroduodenal anastomosis (EGDA) as previously described utilizing male Sprague-Dawley rats (45, 83). Briefly, incisions were made at the gastroesophageal junction (with care not to reach the glandular stomach) and at the anti-mesenteric border of the duodenum, followed by side-to-side duodenoesophageal anastomosis with accurate mucosal to mucosal opposition. The EGDA or anastomosis of the duodenum to the gastroesophageal junction creates chronic reflux of bile and acidic gastric contents into the lower esophagus, mimicking human GERD or reflux-driven BE, the only identified precursor lesion to EAC (84, 85). As shown in Figure 1A, the anastomosis position is critical for inducing reflux of bile mixed with acidic gastric secretions which promotes the development of premalignancy and EAC. Our group observed that surgical failure results if the anastomosis site is not placed at the esophageal gastric junction. Positional surgical failure results in the esophagus being exposed primarily to bile reflux which does not induce premalignancy nor cancer in this model in the absence of acidic gastric excretions.

### Dose range-finding study

To determine the appropriate concentration of C-PAC to utilize in the long-term chemoprevention bioassay we consulted the published literature and performed a 6-week safety and dose-range finding study in Sprague-Dawley rodents (43, 44, 80–82). In the dose-range finding study, rats (n=6/treatment group) received Water or C-PAC in the drinking water at 250, 500 or 700 µg/rat/day *ad libitum*. Water and C-PAC were replenished at least weekly. Blood was drawn in EDTA-treated tubes and centrifuged immediately to permit collection of plasma for serology profiling (Supplemental Figure 1B).

### Long-term chemoprevention bioassay

The study schematic, timeline and experimental groups and are detailed in Figure 1A. The experimental groups included water, C-PAC, reflux, C-PAC+reflux. C-PAC levels were targeted at 350 or 700 µg/rat/day in the drinking water *ad libitum* throughout the study; however, surgical failures in the low dose C-PAC+reflux group resulted in insufficient sample size and exclusion of these animals from the final analysis. The previous week’s water intake was utilized to accurately prepare the targeted concentrations of C-PAC required each week (target: 700 µg/rat/day, actual resultant delivery was 690 µg/rat/day over the course of the 40-week bioassay). Animals were sacrificed at 25 and 40 weeks post-reflux-inducing surgery, and organs were either flash-frozen in liquid nitrogen for molecular analyses or fixed in neutral buffered formalin for 24 h prior to standard processing, paraffin embedding, sectioning and hematoxylin and eosin (H&E) staining for histopathological analysis of tissues. Prior to necropsy, fresh fecal pellets were flash frozen in liquid nitrogen and stored at −80°C until further processing for microbiome analysis.

### Necropsy and histopathology evaluation

At necropsy each rat contributed blood, esophageal and major organ tissues for evaluation. Each esophagus was harvested intact, split longitudinally and characterized for gross changes and the anastomosis site confirmed in terms of position and size opening (≥0.5 cm). Neutral buffered formalin fixed esophagi (6-8 animals per treatment group) were processed at the Children’s Research Institute Histology Core at the Medical College of Wisconsin (Milwaukee, WI). Tissues were paraffin-embedded, 25 5-µm sections prepared, and every fifth section stained with H&E. Histopathological evaluation of full thickness sections included scanning the entire length of each esophagus (50 fields on average) at 100X and blinded grading of each field as normal, hyperplasia, low-grade dysplasia, high-grade dysplasia or EAC.

### Metabolomics and bioinformatic analyses

Sample preparation, extraction, metabolite identification, data handling and initial analysis for untargeted metabolomics were conducted by Metabolon, Inc (Durham, NC). Homogenized esophageal and liver samples were extracted in methanol and dried under vacuum to remove organic solvent. Samples were characterized using Reverse Phase Ultrahigh Performance Liquid Chromatography – Tandem Mass Spectrometry. Compounds were identified based on comparison to a library of authenticated standards maintained by Metabolon, Inc. and statistical analysis conducted in ArrayStudio on log-transformed data. Six samples per treatment group were characterized for metabolomics.

Statistically significant (*P*≤0.05) metabolites induced by reflux and directly reversed by C-PAC (n=200) were uploaded and analyzed in Metacore™ and Cortellis Solution software (https://clarivate.com/products/metacore/, Clarivate Analytics, London, UK). All *P*-value and FDR significant Pathway Maps, Process Networks, Diseases, Go Processes, Metabolic Networks, GO Molecular Functions and GO Localizations for each comparison were exported. Significance was calculated using the hypergeometric test as previously described (86), and it should be noted that Metacore had the highest recognition of metabolites at 86%, which contrasts with reduced metabolite recognition obtained in other programs evaluated including Metaboanalyst (65%), Metscape (45%) or PaintOmics (35%).

Quantitative validation of specific bile acids was conducted at the University of Michigan Metabolomics Core as previously described using liquid chromatography-mass spectrometry separation by reverse phase chromatography (87). Bile acids were measured using negative ionization mode triple quadrupole multiple reaction monitoring methods as previously described (88).

### DNA isolation

Total genomic DNA was isolated from fecal pellets using the PowerLyzer® PowerSoil® DNA Isolation Kit (Mo Bio Laboratories, Carlsbad, CA) according to manufacturer’s instructions with one modification. Fecal samples were transferred to glass bead tubes with 750 µL bead solution, incubated 65°C for 10 min and transferred to 95°C 10 min. DNA samples were eluted with 100 µL of Solution C6 and stored at −80°C until sequencing.

### Microbiome 16S rRNA gene sequencing and bioinformatic analyses

PCR amplicon libraries targeting the 16S rRNA encoding genes present in metagenomic DNA were produced using a barcoded primer set adapted for the Illumina MiSeq (89). DNA sequence data was generated using Illumina paired-end sequencing at the Environmental Sample Preparation and Sequencing Facility at Argonne National Laboratory (Lemont, IL) as previously described (90). Amplicons were sequenced on a 151-bp x 12-bp x 151-bp MiSeq run using customized sequencing primers and procedures (89). Sequence data was processed in QIIME using published protocols (90). Operational taxonomic units (OTU) were identified using Greengenes and aligned with PyNAST to construct phylogenetic trees using Fast Tree 2.0. Plotting and statistical analyses were conducted in R v3.3.0 using phyloseq and EdgeR. There were 5-7 animals per treatment group included in this analysis with sequencing results found in Supplemental Table 16.

Alpha- and beta-diversity were assessed using the Qiagen CLC Genomics Workbench 20.0 with CLC Microbial Genomics Module 20.0 (https://digitalinsights.qiagen.com/). In short, sequencing files were imported into the software, followed by OTU clustering using the default parameters. A phylogenetic tree was reconstructed to perform alpha- and beta-diversity analysis. Alpha diversity was measured using the Chao1 bias-corrected estimation and beta diversity was determined using the Bray-Curtis dissimilarity and Jaccard similarity indexes. The significance of beta-diversity was assessed using the Permutational Multivariate Analysis of Variance (PERMANOVA). The differential abundance analysis tool was used to determine significant changes in OTUs across all samples, and the heatmap was generated using the top 25 most significantly changed OTUs (*P*≤0.05, FDR≤0.05, and Bonferroni-adjusted *P*≤0.05).

### Species level microbiome identification and analysis

The same genomic DNA submitted for 16S rRNA gene sequencing was assessed for species level changes using the Axiom^®^ Microbiome Array (ThermoFisher Scientific, Waltham, MA) on the GeneTitan Multi-channel instrument (ThermoFisher Scientific) at the University of North Carolina at Chapel Hill High-throughput Sequencing facility following manufacturer’s instructions. Data were analyzed using the Axiom Microbial Detection Analysis Software (MiDAS; ThermoFisher Scientific) following manufacturer’s and published (91) protocols to determine the frequency of microbial families and species in each treatment group. Fisher’s exact tests were used to determine significant differences in families or species in gut microbiomes, with tests for trend using the Chi Square test (*P≤*0.05). There were 6-8 animals per treatment group included in this analysis.

### PICRUSt analysis for functional microbial metagenomic profiling

Predicted functions of bacterial communities were identified in PICRUSt based on published methods (92, 93). PICRUSt utilizes existing annotations of gene content and 16S rRNA gene copy number from reference bacterial and archaeal genomes to predict the presence of gene families. Files were formatted as ‘.biom’ and imported into PICRUSt for OTU tables generated using the pick_closed_references_otus script in QIIME (94). The OTU abundances were normalized against the reference 16S rRNA gene copy numbers using the normalize_by_copy_number.py script which divides each OTU by the known or predicted 16S rRNA gene copy number abundance. The normalized table was used to predict functions for the metagenome using the script ‘predict_metagenome.py’. The script outputted a table of function counts as KEGG Ontology (KO) by sample IDs (95, 96). The accuracy of these predictions was evaluated by using the Nearest Sequenced Taxon Index (NSTI). Lastly, predicted KO abundances for the given OTU table were collapsed into individual KEGG pathways by using the script ‘categorize_by_functions.py’. The output files were uploaded to Statistical Analysis of Taxonomic and Functional Profiles (STAMP)^10^, unclassifed reads removed and statistical analysis performed to determine differences between functional pathway data (KEGG pathways Level 2 and 3), as well as 125 identified significant KEGG KOs by ANOVA with Storey’s FDR correction for multiple comparisons and Tukey’s post hoc test. Mean (± standard deviation) Nearest Sequenced Taxon Index (NTSI) values for water, C-PAC, reflux and C-PAC+reflux were 0.11±0.03, 0.12±0.01, 0.10±0.02 and 0.12±0.02, indicating samples were suitable for PICRUSt analysis (92).

### RNA isolation and expression analysis

RNA was isolated from rat lower esophageal tissues using the RNeasy Fibrous Tissue Kit (Qiagen, Germantown, MD). Each sample was homogenized in 400 µL of Buffer RLT with beta-mercaptoethanol for 3 – 10 second pulses with the homogenizer (Pro-Scientific Inc., Oxford, CT) set on level 2. Three to six samples per treatment group were evaluated. RNA was purified following manufacturer’s instructions and eluted in 20 µL of Ambion RNA Storage Solution (Thermo Fisher Scientific, Waltham, MA). RNA concentration and quality was measured using the RNA 6000 Pico kit on the Bioanalyzer 2100 capillary electrophoresis system (Agilent, Santa Clara, CA) and stored at −80°C. One microgram of RNA per sample was reverse transcribed using the iScript™ Advanced cDNA Synthesis Kit (Bio-Rad, Hercules, CA) following manufacturer’s protocol. Expression levels of 85 genes were assessed using the PrimePCR Antibacterial Response (SAB Target List) R384 rat plate (Catalog #10047065; Bio-Rad) using SsoAdvanced Universal SYBR Green Supermix (Bio-Rad). Real-time PCR was performed on the CFX384 real-time PCR system (Bio-Rad) following manufacturer’s protocol. Data were analyzed using CFX Manager (Bio-Rad) where relative changes in gene expression were calculated by 2^-ΔΔCt^, where ΔΔCt = ΔCt (reflux) – ΔCt (water) or ΔΔCt = ΔCt (C-PAC+reflux) – ΔCt (reflux) as previously described (97). Data were normalized to expression levels of *Gapdh* and *Hsp90ab1* with statistical significance determined by Students *T*-test where *P*≤0.05 was considered significant.

### Bacterial response genes identified in human EAC samples

The publicly available GEO dataset (GSE26886), originally contributed by Dr. Wolfgang Kemmner (98), was downloaded from the NCBI Gene Expression Omnibus website. For comparison purposes, Log_2_ transformed data were organized by histopathology and contained a total of 69 human esophageal samples of mixed pathologies. Esophageal squamous cell data (n=9) were removed from our analysis. The remaining data consisted of normal esophageal squamous epithelium (n=19), BE (n=20) and EAC (n=21). Normal epithelium was compared to both BE and EAC. A Students *T*-test with Benjamin-Hochberg FDR correction was utilized to determine statistically significant differences (FDR≤0.05). A second publicly available GEO dataset (GSE193946) containing human tissues from patients with BE with low-grade dysplasia (LGD) to BE with high-grade dysplasia (HGD) and EAC (99) was also utilized to compare gene level changes in the rat esophagus. Results in Table 1 denoted by superscript “a” in column 2 indicate that changes with reflux in the rat esophagus were consistent with significant dysregulation of the same marker in either human GEO dataset.

### Protein isolation and western blot analysis

Rat lower esophageal lysates were prepared by homogenization (PRO Scientific Inc., Oxford, CT) in T-PER® Tissue Protein Extraction Reagent (ThermoFisher Scientific) with cOmplete™ EDTA-free protease inhibitor cocktail and PhosSTOP phosphatase inhibitors (Sigma-Aldrich, St. Louis, MO) according to manufacturer’s instructions. Protein levels in soluble lysates were quantified using the DC protein assay (Bio-Rad) and 15-30 µg/lane loaded into precast 4-20% and 10% Mini-Protean TGX gels (Bio-Rad). Immunoblotting was performed using commercially available antibodies as described in Supplemental Table 17. Images were captured via the ChemiDoc Molecular Imager and band quantification with ImageLab analysis software (both Bio-Rad). Expression values normalized to appropriate loading controls were determined by chemiluminescent immunodetection with fold-change from the Water alone group reported.

### Data integration of metabolites and antimicrobial genes

To build integrated networks, significant antibacterial response genes (*P*-value≤0.05, n=47) and significant metabolites (*P*-value≤0.05) for reflux vs water (n=319) and C-PAC+reflux vs reflux (n=264) were uploaded individually into Metacore™ and Cortellis Solution software (Clarivate Analytics, London, UK). The C-PAC+reflux vs reflux gene and metabolite lists were activated and the analyze network feature applied to build networks with 100 nodes using all compound-target interactions. The “analyze network” feature first builds one large network, and this network is then divided into smaller fragments based on the chosen node size for easier visualization and exploration. The result is a list of 30 partially overlapping networks, each with associated GO-Processes, P-values, g-scores and z-scores calculated as previously described (86) and select networks were visualized.

## Statistics

Additional statistical analyses not described elsewhere were performed in GraphPad Prism software (La Jolla, CA). Body weight, food consumption and water consumption data were analyzed for differences between all treatment groups using repeated measures ANOVA where time was considered a variable and with Tukey’s post-hoc test. All *P*-values ≤ 0.05 were considered significant.

## Study approval

All experimental procedures were conducted in accordance with a protocol approved by the Institutional Laboratory Animal Care and Use Committee at the Medical College of Wisconsin (AUA3095) and consistent with the National Institutes of Health Guide for the Care and Use of Laboratory Animals.

## Authors’ contributions

LAK and KW conceived and designed the experiments. LAK, KMW and CLH collected and/or contributed data. ABH prepared and provided cranberry proanthocyanidins. LAK, KMW, CLH, YZ, JHR, JAA, MW, BAT and JLC performed data analyses. LAK and KMW wrote the paper assisted by CLH, YZ, BAT, JLC, ABH, JHR, JAA and MW. All authors reviewed and approved of the final manuscript.

## Acknowledgments

We thank the National Cancer Institute (R01CA158319), the University of Michigan (U057239) and the John and Carla Klein family research fund, all awarded to Laura A. Kresty for supporting this study. Additional support was provided through U54CA163059. Dr. Abrams was supported in part by the National Cancer Institute (U54CA163004, R01CA238433, R01CA255298, R01CA272898). We would like to thank Dr. Nita Salzman and Dr. Meredith Halling at the Medical College of Wisconsin for their expertise in microbiome isolation techniques and rodent surgeries, respectively. Figure 8 was generated using BioRender (BioRender.com).

